# Liver-to-Atria Inflammatory Axis Driving Arrhythmia

**DOI:** 10.64898/2026.05.19.726408

**Authors:** Yue Yuan, Shuyue Wang, Jifei Ding, Jun Jiang, Yuying Zeng, Tingting Li, Anna Kay Shinohara, Chunhao Lin, Carole W. Sun, Ron Cornelis Hoogeveen, Mihail G. Chelu, Seyedmohammad Saadatagah, Sung Yun Jung, Danyvid Olivares-Villagómez, Christie M. Ballantyne, Bingning Dong, Na Li

**Affiliations:** Department of Medicine, Section of Cardiovascular Research, Baylor College of Medicine, Houston, Texas, USA; Department of Medicine, Section of Gastroenterology and Hepatology, Baylor College of Medicine, Houston, Texas, USA; Department of Medicine, Section of Cardiology, Baylor College of Medicine, Houston, Texas, USA; Center for Translational Research on Inflammatory Diseases, Baylor College of Medicine, Houston, Texas, USA; Department of Biochemistry and Molecular Pharmacology, Baylor College of Medicine, Houston, Texas, USA; Division of Infectious Diseases, Department of Medicine, Vanderbilt University Medical Center, Nashville, Tennessee, USA

**Author notes:** Corresponding author: Bingning Dong, Ph.D., Department of Medicine, Baylor College of Medicine, 1 Baylor Plaza, BCM23, Houston, Texas, 77030, USA, Na Li, Ph.D., Department of Medicine, Baylor College of Medicine, 1 Baylor Plaza, BCM285, Houston, Texas, 77030, USA. These authors contributed equally to this work (Yue Yuan, Shuyue Wang). These authors contributed equally to this work (Bingning Dong, Na Li). **Competing interests:** All authors declare that there are no competing interests.

**Keywords:** Metabolic dysfunction–associated steatohepatitis, atrial fibrillation, osteopontin, SPP1, gasdermin D

## Abstract

**Background:** Metabolic dysfunction–associated steatohepatitis (MASH) is emerging as a risk factor of cardiometabolic diseases, including the atrial fibrillation (AF) - the most common sustained arrhythmia. Given that the liver is a major source of inflammatory mediators, lipids, and hepatokines under metabolic stress, we hypothesized that hepatocyte-derived factors in MASH may accelerate atrial remodeling and arrhythmogenesis.

**Methods:** Analysis of the Atherosclerosis Risk in Communities (ARIC) visit 5 cohort was performed to determine the association between the FIB-4 index - a classic indicator of liver fibrosis, and AF risk, with multivariable adjustment for common comorbidities. A murine model of MASH was induced using the GAN (Gubra-Amylin NASH) diet. Programmed intracardiac stimulation and echocardiography were performed to assess AF susceptibility and cardiac function. Calcium imaging, histology, flow cytometry, plasma proteomics, and single-nucleus RNA sequencing (snRNA-seq) analyses were employed to elucidate the role of recruited inflammatory macrophages via hepatocyte-derived osteopontin (OPN) in MASH-induced atrial remodeling.

**Results:** Analysis of the ARIC cohort confirmed a higher cumulative incidence of AF and an elevated adjusted hazard ratio (HR) in patients with intermediate and high FIB-4 indices compared to individuals with low FIB-4 scores. MASH mice exhibited increased susceptibility to pacing-induced AF, accompanied by enhanced proarrhythmic calcium release events, atrial enlargement, and fibrosis, independent of ventricular dysfunction. Proteomics and snRNA-seq revealed that the hepatocyte-secreted OPN under MASH conditions promoted the differentiation and recruitment of TGFBR1^+^ inflammatory macrophages to the atria, leading to gasdermin D (GSDMD) activation – an effector of inflammasome signaling and consequent proarrhythmic atrial remodeling. Activation of the monocyte-derived pro-inflammatory TGFBR1^+^ macrophages was dependent on the OPN receptor CD44. Furthermore, the MASH-induced atrial fibroinflammatory milieu and enhanced AF susceptibility were mitigated through several strategies, including hepatocyte-specific *Spp1* (encoding OPN) deletion, neutralization of circulating OPN, ablation of CD44 or GSDMD.

**Conclusions:** These findings establish a pathogenic role of the hepatokine osteopontin in driving activation and recruitment of TGFBR1^+^ inflammatory macrophages into the atria, leading to proarrhythmic atrial remodeling under MASH. Osteopontin-targeted therapy or GSDMD inhibition prevents AF, indicating a novel therapeutic strategy for liver disease-related atrial arrhythmogenesis.

**Clinical Perspective:** *What is new?:*

- In the ARIC cohort, metabolic dysfunction-associated steatohepatitis (MASH) is associated with increased risk of atrial fibrillation (AF) after adjusting for common comorbidities. Elevated levels of circulating osteopontin (encoded by SPP1) predict an increased risk of AF in patients with MASH-induced liver fibrosis.
- MASH enhances hepatocyte secretion of osteopontin, leading to expansion of myeloid cells and recruitment of inflammatory macrophages into atria. This liver-to-atrial inflammatory circuit promotes the development of a substrate conducive to AF, which can be attenuated by hepatocyte-specific Spp1 deletion or neutralizing anti-anti-osteopontin antibody treatment to eliminate the mediator, or ablation of inflammasome effector gasdermin D to correct the atrial response.

*What are the clinical implications?:*

- Osteopontin may serve as a biomarker for AF in MASH cohorts.
- Anti-osteopontin therapy through neutralizing antibodies may serve as a novel therapeutic strategy for liver disease-related atrial arrhythmia.

## Introduction

Atrial fibrillation (AF) is the most common sustained cardiac arrhythmia with severe complications ^1,2^. Although rhythm control remains the cornerstone^3,4^, upstream strategies that target causal risk factors and disease substrates are needed to improve long-term AF management. Metabolic dysfunction-associated steatotic liver disease (MASLD) affects nearly one-third of the adult population worldwide^5,6^. Its progressive form, metabolic dysfunction-associated steatohepatitis (MASH), is characterized by hepatic inflammation and cellular injury that can progress to fibrosis, cirrhosis, and hepatocellular carcinoma^7^. Patients with MASH have increased risk for all-cause mortality with cardiovascular disease (CVD) as a leading cause of death^8^. Epidemiological studies link MASLD/MASH to higher AF incidence^9,10^; however, whether MASH directly contributes to AF pathogenesis, and through which mechanisms, remains unclear. Intriguingly, both AF and MASH share convergent pathophysiological features: chronic inflammation, metabolic disturbance, and fibrosis, suggesting they may be driven by a common fibroinflammatory signal. Infiltrating inflammatory monocytes promote proarrhythmogenic tissue remodeling by interacting with resident fibroblasts, increasing conduction heterogeneity and AF predisposition^11–13^. Given that the liver is a major source of inflammatory mediators, lipids, and hepatokines under metabolic stress, we hypothesized that hepatocyte-derived factors in MASH may accelerate atrial remodeling and arrhythmogenesis.

Here, we demonstrate that MASH progression enhances AF susceptibility by promoting inflammatory macrophages expansion, disrupting myocardial calcium handling, and promoting atrial fibrosis. Hepatocyte-secreted osteopontin (OPN), encoded by *SPP1*, predicts AF risk in MASH patients with advanced liver fibrosis. In a murine model, osteopontin mediates MASH-induced arrhythmogenesis through activation and recruitment of pro-inflammatory and pro-fibrotic macrophages, an effect that can be mitigated by hepatocyte-specific *Spp1* knockout, neutralizing anti-OPN antibodies, ablation of CD44 (a key receptor of OPN on macrophages), or ablation of inflammasome effector gasdermin D (GSDMD). This study uncovers a molecular pathway underlying the liver-to-atria inflammatory circuit that drives arrhythmogenesis.

## Methods

Detailed methods are provided in the Supplemental Material. All data are available in the main text or the supplementary materials.

### Epidemiological studies

Human data were obtained from the Atherosclerosis Risk in Communities (ARIC) study, a prospective population-based study. At baseline men and women, aged 45 – 64, were recruited from 4 communities in the US: Forsyth Co, NC; Jackson, MS; northwest suburbs of Minneapolis, MN; and Washington County, MD. For the current analysis, we included individuals attending Visit 5 (n=6,538) conducted between 2011 and 2013 and follow-up through Dec 2021. We excluded participants with chronic liver diseases such as HBV/HCV-related hepatitis, missing data on liver enzymes or platelet counts (n=247), race other than African American or whites (n=18), non-whites in the Minneapolis and Washington County field centers due to low recruitment numbers (n=20), excessive alcohol intake (defined as ≥196 g usual ethanol intake per week for men and ≥98 g usual ethanol intake per week for women^14,15^) or missing information on alcohol intake (n=722) (**Supplemental Figure S1**). For the prospective analyses, we also excluded participants with AF at baseline. The study protocol was approved by the institutional review boards of all participating centers, and all participants provided written informed consent. Fibrosis-4 (FIB-4) index, calculated as FIB-4 = age (years) × AST(U/L) / (platelet count [10^9^/L] × ALT [U/L]^1/2^), is a non-invasive score for advanced liver fibrosis and was used as a surrogate for MASH. Given the advanced age of enrolled participants, age-adjusted FIB-4 cutoffs were applied to classify the degree of liver fibrosis: <1.45 (low), 1.45 – 3.25 (intermediate), and >3.25 (high), as previously described^16^. The primary outcome was incident AF. In the ARIC cohort, AF events were identified using electrocardiogram readings obtained during visits, hospital discharge diagnoses, and death certificates and were validated by an expert committee. Osteopontin levels were measured among 5000 plasma proteins using a multiplexed modified DNA-based aptamer (SomaScan) technology. Briefly, a slow off-rate modified aptamer (SOMAmer) reagents (SomaLogic, Inc, Boulder, Colorado) capture proteins from blood samples, then the SOMAmer reagents were measured in fluorescent arrays. The relative concentration of proteins was then derived from the concentration of SOMAmer reagents. All procedures performed in studies involving human participants were in accordance with the institutional and national research committee and the Declaration of Helsinki.

### Animal studies

All procedures for mouse experiments were approved by the Institutional Animal Care and Use Committee at Baylor College of Medicine (Protocol number AN-7259 and AN-1550) and confirmed to the Guide for the Care and Use of Laboratory Animals published by National Institutes of Health. Mice were housed in groups of 2-5 animals in IVC cages under a strict 12:12 h light/dark cycle at central animal facility of Baylor College of Medicine. All mice were free to get food and water. Both male and female mice were included in this study. To establish MASH in mice, C57BL/6J wildtype (WT) mice of 2-month-old were randomly subjected to Gubra-amylin NASH (GAN) diet (including 40 kcal% fat, 30% fructose, 10% sucrose and 2% cholesterol, Research diets, Inc. Cat# D09100310) for 5 months. Chow diet-fed WT mice served as controls. To elucidate molecular mechanisms, multiple genetic knockout mouse lines were established including hepatocyte-specific *Spp1* knockout mice^17^ (*Spp1^f/f^*;*Alb^Cre^*), CD44 whole body knockout mice (*Cd44^-/-^*), GSDMD whole body knockout mice (*Gsdmd^-/-^*), and cardiomyocytes-specific GSDMD knockout mice (*Gsdmd^f/f^*;*Myh6^CreER^*). All strains were on a C57BL/6J background. For intervention studies, 2-month-old mice were subjected to the GAN diet for 4 months initially, then randomized to receive the treatment of anti-OPN neutralizing antibody (100µg per mouse per dose, twice weekly, i.p. injections) or IgG antibody during the last month of the GAN diet. The anti-OPN neutralizing antibodies and IgG1 isotype control were purchased from BioXcell (Cat# BE0382, BE0083). Resmetirom was purchased from Ambeed (Cat# AMBH303C650B).

### Statistical analysis

Baseline characteristics of human data were tabulated by FIB-4 status, with continuous variables reported as mean (SD) or median (IQR) and categorical variables as number (percentage). The association of exposures with incident AF was studied using Cox proportional hazards models in a stepwise adjustment fashion. P trend for linearity of hazard ratio of proportional hazard regression model was calculated based on the results of Wald chi-square test on linearity hypothesis of ordered MASH likelihood categories. Given the advanced age of participants from the ARIC cohort and the high mortality, we performed a sensitivity analysis to consider the competing risk of death. For animal studies, normality was performed with Anderson-Darling test or Shapiro-Wilk test. All mouse data were presented as mean ± SEM or percentage where applicable. Statistical analyses were performed using GraphPad Prism 10.1. Two-tailed Student’s *t*-tests were used to compare data between two groups maintaining normal distributions. Mann-Whitney tests were used for data where normality could not be assumed. Fisher’s exact test was used to analyze categorical data. Nested t-tests or RStudio were used for data of biological replicates with repeated measurements. Multilevel models were implemented in RStudio using lme4, and P-values were derived employing the Kenward-Roger approximation. A P-value less than 0.05 was considered statistically significant.

## Results

### MASH promotes atrial arrhythmogenesis

To determine the relation between the severity of MASH-associated fibrosis and AF incidence in patients, we analyzed the human data from the ARIC study visit 5 conducted between 2011 and 2013 and followed up through 2021. The FIB-4 index was used as a surrogate of MASH-associated liver fibrosis^18–20^. After exclusion of those with heavy drinking status, 5,051 participants (mean age of 76 (SD 5.2) years, 58% female, 22% black) were included (**Supplemental Figure S1, Table S1**). After adjusting for multiple common comorbidities including age, sex, BMI, cholesterol, diabetes, hypertension, and triglycerides (Model 3), during a mean follow-up of 7.6 ± 2.78 years, the cumulative incidence of AF and adjusted hazard ratio (HR) were greater in patients with intermediate (n=2,931, 19.14%, [HR: 1.20, 95%CI: 1.02 - 1.40]) and high FIB-4 (n=260, 24.62%, [HR: 1.51, 95%CI: 1.13 - 2.03]) indices compared to patients with low FIB-4 index (n=1,860, 13.76%, ref) (**Figure 1A; Supplemental Table S2**), and remained significant or trending toward significance after excluding advanced kidney disease (eGFR<30, **Supplemental Table S3**) or excluding prevalent heart failure (HF) (**Supplemental Table S4**).

**Figure 1.**
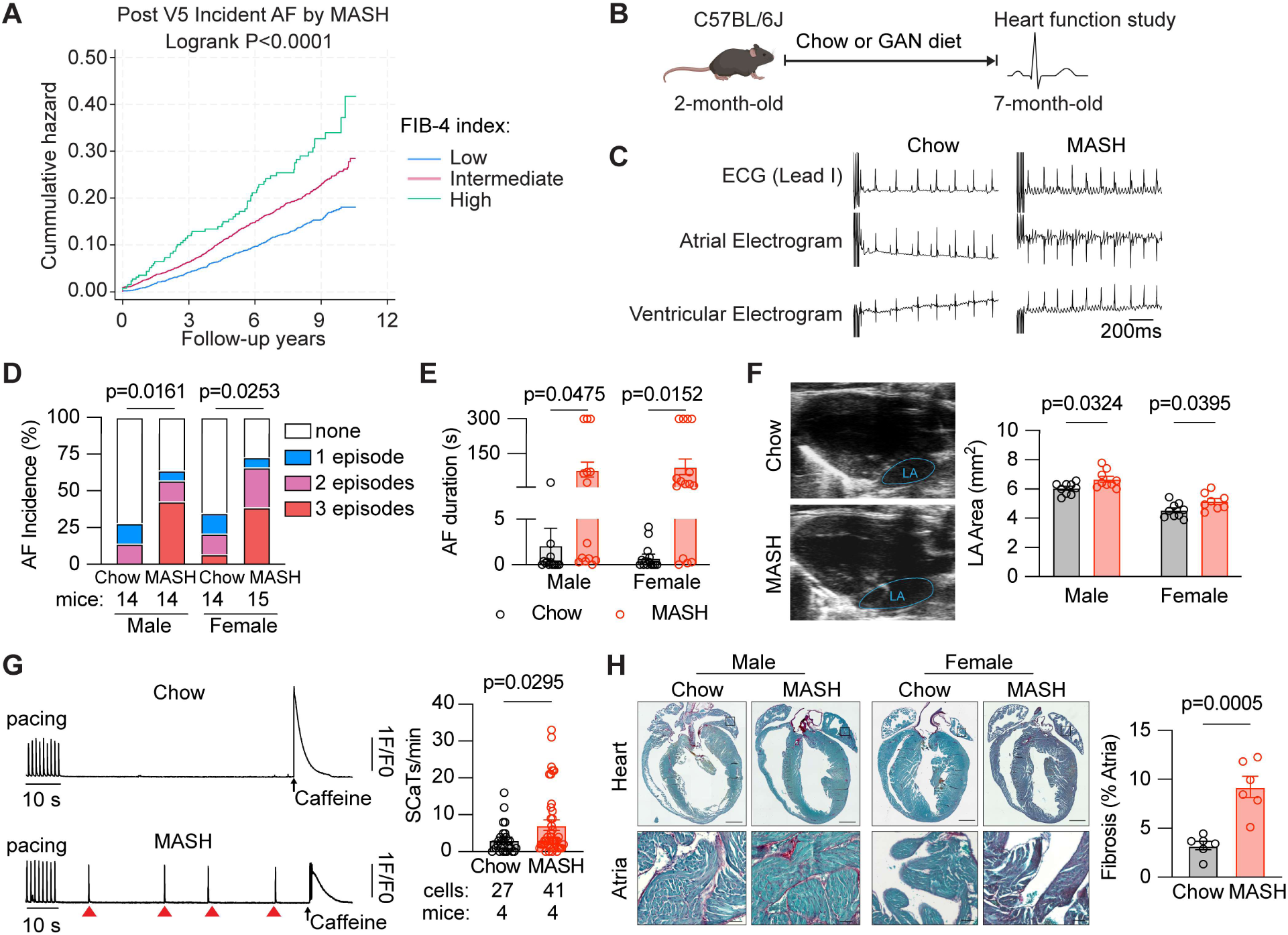
MASH promotes atrial arrhythmogenesis. (**A**) ARIC Post Visit 5 Aalen–Nelson cumulative hazard curves for incident AF events stratified by FIB-4 categories over a mean follow-up of 7.6 years. (**B**) Schematic illustrating the 5-month GAN diet-induced MASH model. (**C**) Representative surface ECG and intracardiac electrograms in Chow and MASH mice. (**D-E**) Incidence (**D**) and duration (**E**) of pacing-induced AF incidence. (**F**) Representative B-Mode echocardiography of Chow and MASH mice, and measurements of LA area in Chow and MASH mice, n=8 or 9 per group. (**G**) Representative traces of Ca^2+^ transients induced by 1-Hz pacing in isolated atrial cardiomyocytes, followed by baseline recording and acute application of 10 mmol/L caffeine. The frequency of spontaneous Ca^2+^ transients (SCaTs, red arrows) in Chow and MASH mice. (**H**) Representative Picro-sirius staining of four-chamber heart in Chow and MASH mice. Quantification of atrial fibrosis levels in Chow and MASH mice, n=3 male and 3 female mice per group. The bar graph data are mean ± SEM with individual values. p-values are determined with two-tailed Student’s t-test in **E**, **F, G,** and **H**; Mann–Whitney test in **D** .

To investigate the mechanistic basis of this association, we used a murine model of diet induced MASH. Two-month-old C57BL/6J mice were fed either normal chow or GAN diet, respectively (**Figure 1B**). After 5 months of GAN feeding, mice developed typical characteristics of MASH, including increased body weight (BW), liver weight, liver-to-body weight ratio, fasting plasma glucose (FPG) levels, plasma and liver triglyceride (TG), and total cholesterol (TC) levels, as well as serum alanine aminotransferase (ALT) levels, compared with chow-fed controls (**Supplemental Figure S2A-E**). Histological studies revealed worsened steatosis, inflammatory foci, collagen deposition, and F4/80^+^ macrophage accumulation in the livers of MASH mice. Consistently, expression of lipid metabolism (*Srebp1*, *Fasn, Scd1,* and *Pparα*), inflammation (*F4/80, Tnf, Mcp1,* and *Il1b*), and fibrosis (*Acta2*, *Co1a1,* and *Tgfb*) markers were markedly upregulated in the livers of MASH compared to chow mice (**Supplemental Figure S2F, G**). At this stage of established MASH, surface ECG recording revealed the development of short episodes of spontaneous AF that self-terminated within a few seconds in MASH mice (**Supplemental Figure S3A-C**). To assess whether MASH mice developed a substrate for AF, programmed electrical stimulation (PES) study was conducted to evoke AF. MASH mice developed both higher incidence and longer duration of reproducible pacing-induced AF episodes compared with chow-fed mice (**Figure 1C-E**). The left atria were enlarged in MASH mice (**Figure 1F**), whereas blood pressure, left ventricular systolic and diastolic functions, and exercise tolerance were comparable to those of the chow group (**Supplemental Figure S2D-G**). These data suggest that MASH induces a pro-arrhythmic substrate independent of ventricular dysfunction. Consistent with echocardiographic data, histological analysis demonstrated larger atrial size in the MASH group, accompanied by increased atrial weight (**Supplemental Figure S2H,I**). Given that male and female MASH mice exhibited similar degrees of liver phenotype and AF susceptibility, we combined data from both sexes in the following figures. The increased AF vulnerability in MASH model was associated with more frequent spontaneous Ca^2+^ transients (SCaTs) in isolated atrial cardiomyocytes (**Figure 1G**), indicating aberrant Ca^2+^ release that contributes to ectopic activities under MASH conditions. The MASH model also exhibited increased collagen deposition and elevated protein levels of fibrotic markers such as collagen A1 (COLA1), matrix metalloproteinase-9 (MMP9), and vimentin (**Figure 1H; Supplemental Figure S2J**). Collectively, these findings indicate that MASH promotes AF development through ectopic activities and structural remodeling.

### Hepatic osteopontin drives MASH-associated atrial arrhythmogenesis

Hepatokines mediate immune activation and cell recruitment to distant tissues under stress ^21^. Osteopontin (OPN, encoded by *SPP1*), a hepatokine released by injured hepatocytes, promotes a systemic inflammatory response and liver fibrosis during MASH progression^22^. To explore whether circulating OPN levels correlate with incident AF, we stratified ARIC participants based on their OPN levels measured on the SomaScan platform. The incidence of AF was greater in those with higher levels of OPN (**Figure 2A**; **Supplemental Figure S4A, Tables S5, S6, S7**). The association between FIB-4 and AF was only statistically significant in those with high OPN levels (**Supplemental Tables S8, S9, S10**). Intriguingly, plasma proteomic profiling in MASH mice also identified OPN among the most significantly elevated proteins (**Figure 2B**; **Supplemental Figure S4B**). OPN protein and *Spp1* mRNA levels were increased in the livers, but not in kidneys or atrial tissues, of MASH mice (**Supplemental Figure S4C-F**). To determine whether hepatocytes are a major source of OPN within the liver, we analyzed publicly available single-nucleus RNA sequencing (snRNA-seq) datasets of liver samples from humans (GES256398)^23^ and a mouse model (GES262939)^24^ of MASH. We found upregulation of *Spp1* in subsets of hepatocytes of both human and mouse MASH samples compared with healthy controls (**Supplemental Figure S4G-H**). Consistently, *Spp1* mRNA levels were upregulated in the isolated hepatocytes from MASH mice (**Figure 2C**). OPN levels were elevated in the conditioned medium of primary hepatocytes, but not Kupffer cells (hepatic macrophages), isolated from MASH mice compared with chow-fed controls (**Figure 2D**). These results indicate that stressed hepatocytes could be a major source of liver-derived osteopontin under MASH conditions.

**Figure 2.**
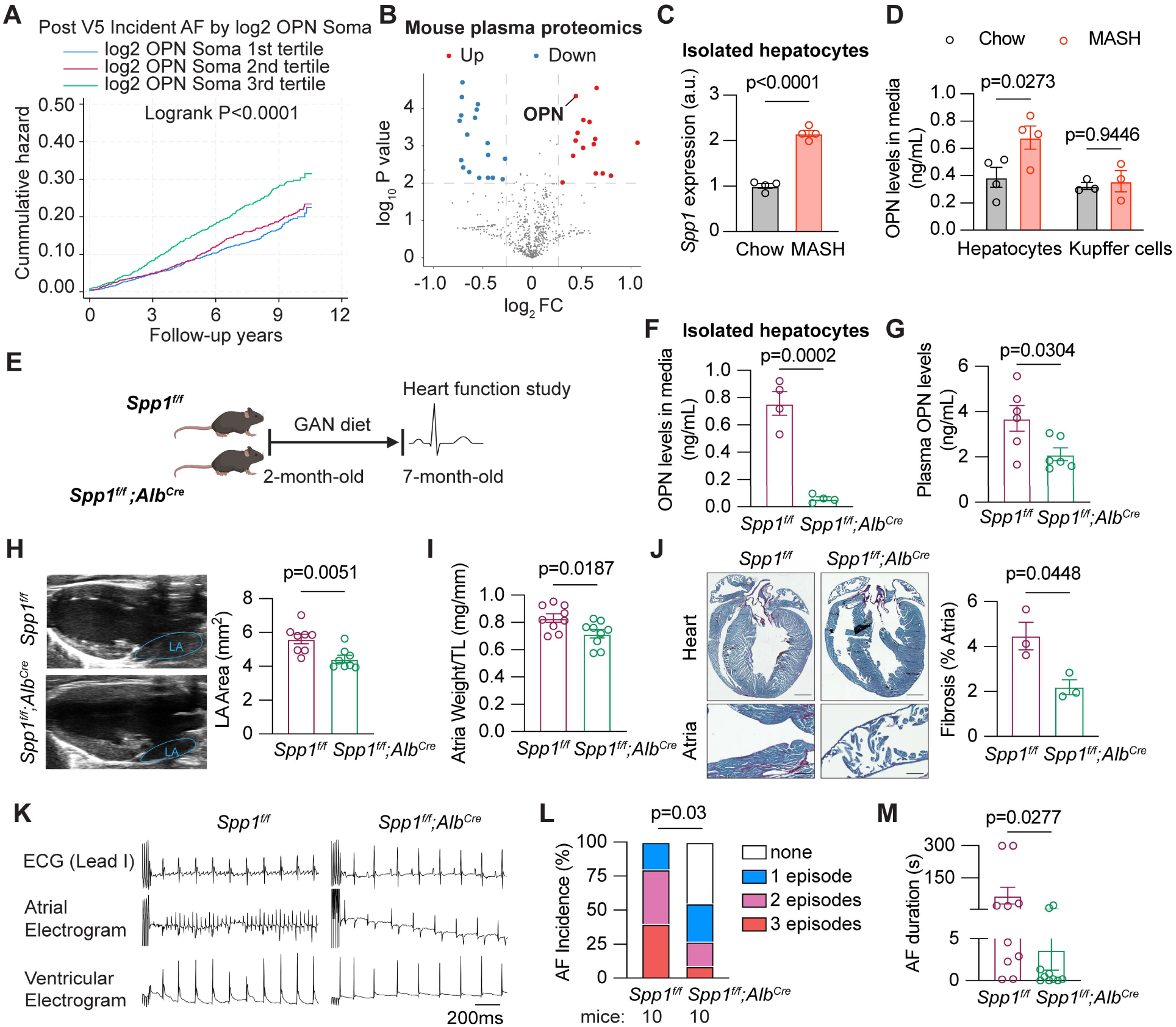
Hepatic osteopontin (OPN) drives MASH-associated atrial arrhythmogenesis. (**A**) ARIC Post Visit 5 Aalen-Nelson cumulative hazard estimation of curve of incident AF event across log2 OPN Soma tertile categories over a mean follow-up of 7.6 years. (**B**) Volcano plot showing differential plasma proteins between Chow and MASH mice, n=3 per group. (**C**) *Spp1* mRNA levels in isolated hepatocytes, n=3 per group. (**D**) OPN levels in cultured conditioned medium of primary hepatocytes or Kupffer cells, n= 4 or 5 per group. (**E**) Scheme of *Spp1^f/f^* or *Spp1^f/f^;Alb^Cre^*mice fed with GAN diet. (**F**) OPN levels in conditioned medium of primary hepatocytes, n=4 per group. (**G**) Plasma levels of OPN, n = 6 per group. (**H**) Representative B-Mode echocardiography images and measurements of LA area in *Spp1^f/f^* and *Spp1^f/f^;Alb^Cre^* mice fed with GAN diet, n=8 per group. (**I**) Quantification of atrial weight to tibial length (TL), n= 9 per group. (**J**) Representative Picro-sirius staining images of four-chamber heart and quantification of atrial fibrosis, n=3 per group. (**K**) Representative surface ECG and intracardiac electrograms in *Spp1^f/f^* and *Spp1^f/f^;Alb^Cre^* mice fed with GAN diet. (**L-M**) Incidence (**L**) and duration (**M**) of pacing-induced AF in two groups. The bar graph data are as mean ± SEM with individual values. p-values are determined with two-tailed Student’s t-test in **C, D, F, G, H, I, J** and **M**; Mann–Whitney test in **L**.

Because a previous study reported that *Spp1*^high^ monocytes can infiltrate atria and promote fibrotic proarrhythmic remodeling in a ‘triple-hit’ model ^25^, we hypothesized that MASH-induced atrial arrhythmogenesis is mediated by circulating OPN derived from injured hepatocytes. To test this, we subjected the hepatocyte-specific *Spp1* knockout mice (*Spp1^f/f^;Alb^Cre^*) and *Spp1^f/f^* controls to GAN diet for 5 months (**Figure 2E**). Western blots confirmed near-complete deletion of OPN protein in isolated hepatocytes of *Spp1^f/f^;Alb^Cre^* mice compared with *Spp1^f/f^*controls (**Supplemental Figure S5A**). Deletion of *Spp1* significantly reduced the OPN levels in the hepatocyte-conditioned medium and plasma of *Spp1^f/f^;Alb^Cre^* mice fed with GAN diet (**Figure 2F,G**). *Spp1^f/f^;Alb^Cre^* mice also exhibited improved liver metabolism and alleviation of liver steatosis, inflammation, and fibrosis compared with *Spp1^f/f^* controls fed with GAN diet (**Supplemental Figure S5B-G**). Atrial structural remodeling and fibrosis were attenuated in *Spp1^f/f^;Alb^Cre^* GAN group compared with *Spp1^f/f^* GAN group, without affecting ventricular function (**Figure 2H-J; Supplemental Figure S5H, Table S11**). Ultimately, the incidence and duration of pacing-induced AF were significantly reduced in *Spp1^f/f^;Alb^Cre^* mice compared with *Spp1^f/f^* controls (**Figure 2K-M**). These findings suggest that hepatocyte-derived osteopontin is a key hepatokine that mediates MASH-associated atrial arrhythmogenesis.

### Hepatic osteopontin promotes a fibroinflammatory milieu in the atria under MASH

Inflammatory remodeling is a common pathophysiological feature in MASH and atrial arrhythmia^26^. Flow cytometry and ELISA revealed the expansion of circulating myeloid cells and Ly6C^high^ monocytes, and elevated inflammatory cytokines levels including IL-1β, IL-6, and TNF-α in MASH mice (**Supplemental Figure S6A-C**). Interestingly, compared with *Spp1^f/f^* GAN mice, deletion of *Spp1* in hepatocytes reduced the numbers of circulating myeloid cells and Ly6C^high^ monocytes, as well as plasma IL-1β and IL-6 levels in *Spp1^f/f^;Alb^Cre^* GAN mice (**Supplemental Figure S6D, E**), suggesting that hepatocyte-derived osteopontin contributes to systemic pro-inflammatory responses associated with MASH.

To further elucidate the atrial milieu under MASH, we performed snRNA-seq on atrial tissues from four groups of mice: Chow, MASH, *Spp1^f/f^* GAN, and *Spp1^f/f^;Alb^Cre^* GAN mice. After quality control and filtering, a total of 121,975 single nuclei were profiled (**Supplemental Figure S7A-E**). Twelve distinct cell populations were identified based on canonical marker genes, including atrial cardiomyocytes (ACMs), endothelial cells (ECs), fibroblasts (FBs), mononuclear phagocytes and dendritic cells (MP/DCs), lymphatic endothelial cells (LECs), and others (**Figure 3A**; **Supplemental Figure S6F**). Compared with the chow group, MASH mice displayed a twofold MP/DC expansion and fewer ACMs in atria; depletion of hepatocyte *Spp1* partially reversed MP/DC and EC proportion in *Spp1^f/f^;Alb^Cre^* GAN group compared with *Spp1^f/f^* GAN group (**Figure 3B**). Gene Ontology (GO) of pseudo-bulk expression revealed the top 5 upregulated pathways related to enhanced immune responses, while the top 5 downregulated pathways associated with cardiac electrical or contractile programs in MASH atria (**Figure 3C**), consistent with the electrical remodeling described earlier. Milo testing identified differentially abundant neighborhoods, with MP/DC, FB, and EC clusters exhibiting most dynamically altered subsets in MASH atria compared with chow controls. In contrast, these altered neighborhoods within MP/DC, FB, and EC clusters were mostly reversed by hepatocyte-deletion of *Spp1* (**Figure 3D**). Specifically, MP/DCs were hyperactive in MASH, with upregulation of cytokine production pathways such IL-6 and IL-1, and canonical inflammasome pathway genes, *Ccr2*, *Trem2*, and *Tgfbr1*; whereas deletion of hepatocyte *Spp1* dampened pathways related to protein ubiquitination, endoplasmic reticulum stress, leukocyte activation, as well as inflammasome pathway-related genes within the MP/DC cluster (**Figure 3E-F**).

**Figure 3.**
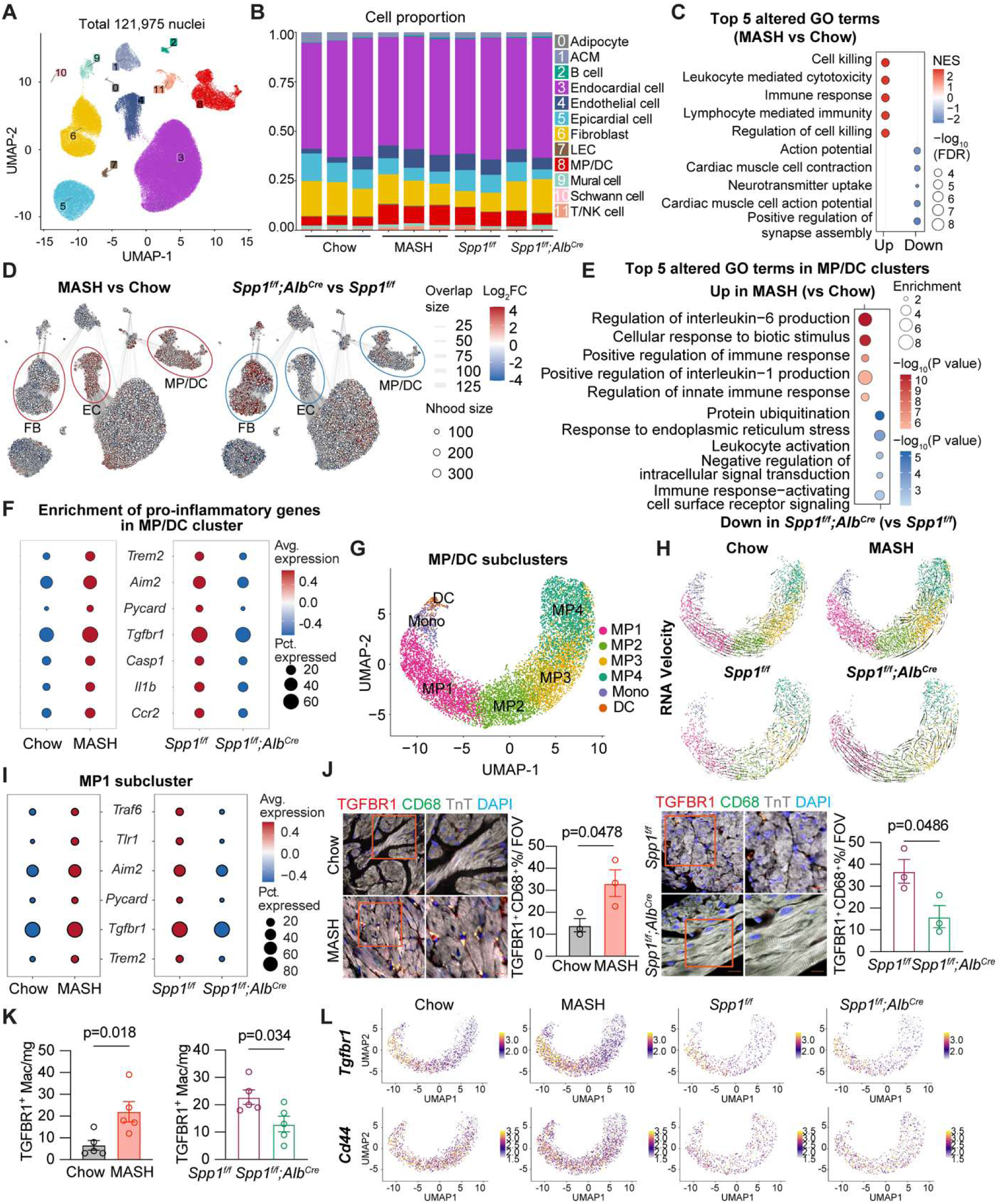
MASH promotes a fibroinflammatory milieu in atria. (**A**) Uniform Manifold Approximation and Projection (UMAP) visualization of 12 distinct cell populations. (**B**) Cell proportion in each sequencing sample. n=2-3 sequencing samples per group. (**C**) GO analysis of the top five upregulated and downregulated pathways based on pseudo-bulk DEGs. (**D**) Milo analysis showing differential cell abundance in each cell population. The MP/DC, FB, and EC clusters were in circles. (**E**) GO analysis of the top five upregulated (MASH vs Chow) and downregulated (*Spp1^f/f^;Alb^Cre^* vs *Spp1^f/f^*) pathways in the MP/DC cluster. (**F**) Dotplot of differential expressions of pro-inflammatory genes in MP/DC cluster among paired comparisons. (**G**) UMAP visualization of MP/DC subsets. (**H**) RNA velocity of MP/DC subsets in Chow, MASH, *Spp1^f/f^* GAN, and *Spp1^f/f^;Alb^Cre^* GAN groups. (**F**) Dotplot of differential expressions of pro-inflammatory genes in MP1 cluster among paired comparisons. (**J**) Representative images of MP1 in mouse atria. Scale bars, 20 or 10 μm. n=3 mice per group. (**K**) Flow cytometry analysis of TGFBR1^+^ proinflammatory macrophages in mouse atrial tissue. n=5 mice per group. (**L**) *Tgfbr1* and *Cd44* expression patterns across MP/DC subsets in 4 groups of samples. Bar graph data are as mean ± SEM with individual values. p-values are determined with two-tailed Student’s t-test in **J** and **K**.

### Hepatic osteopontin enhances expansion of proinflammatory *Tgfbr1*^high^ macrophages under MASH

Within the MP/DC cluster, we identified Mono, DC, and 4 MP subclusters, with Mono, MP1 and MP4 subsets exhibiting expansion in MASH atria compared with chow controls (**Figure 3G**; **Supplemental Figure S7G,H**). MP1 subset expressed high levels of *Tgfbr1* and antigen-representation genes (*H2-Ab1*, *H2-Aa*), indicative of its proinflammatory status (**Figure 3I**, **Supplemental Figure S7G**). RNA velocity analysis revealed that the MP1 cells were primarily originated from proinflammatory Mono cells and other MP subsets, and this transition was enhanced in MASH compared with chow controls. Importantly, deletion of hepatocyte *Spp1* curtailed MP1 transitions from Mono and other MP subsets (**Figure 3H**). Moreover, innate inflammatory pathway-related genes (*Traf6, Tlr1, Aim2, Pycard, and Trem2*) were upregulated in the MP1 cluster of MASH mice compared with chow controls, and these changes were attenuated in *Spp1^f/f^;Alb^Cre^* GAN group compared with *Spp1^f/f^* GAN group (**Figure 3I**). Consistent with the snRNA-seq data, immunofluorescence and flow cytometry confirmed increased TGFBR1^+^ inflammatory macrophages in MASH atria relative to chow controls, whereas *Spp1^f/f^;Alb^Cre^* GAN mice showed markedly reduced TGFBR1^+^ inflammatory macrophages compared with *Spp1^f/f^*GAN mice (**Figure 3J, K; Supplemental Figure S7I**). These findings suggest that MASH promotes proinflammatory *Tgfbr1*^high^ macrophages via the hepatocyte-derived osteopontin.

CD44 is a known osteopontin receptor mediating leukocyte recruitment^27^. In MASH atria, *Cd44,* along with *Tgfbr1*, was upregulated in MP1 cells (**Figure 3L**). Interestingly, not only were CD44 protein levels were increased in bone marrow cells (BMs), but the peripheral CCR2^+^TGFBR1^+^CD44^+^ proinflammatory monocytes also expanded in MASH mice compared with controls (**Supplemental Figure S7A, B**). In contrast, deletion of hepatocyte *Spp1* negatively impacted CD44 protein expression in BMs and the circulating CCR2^+^TGFBR1^+^CD44^+^ proinflammatory monocytes in *Spp1^f/f^;Alb^Cre^* GAN mice compared with *Spp1^f/f^* GAN mice (**Supplemental Figure S7C, D**). To clarify the relation between hepatocyte derived OPN and TGFBR1^+^CD44^+^ macrophage activation, we subjected the primary bone marrow-derived macrophages (BMDMs) to conditioned medium (CM) derived from hepatocytes isolated from chow controls, MASH, *Spp1^f/^*^f^ GAN, and *Spp1^f/f^;Alb^Cre^*GAN mice (**Figure 4A**). Western blots revealed that the hepatocyte-CM from MASH (versus control CM) increased protein levels of TGFBR1 and CD44 in BMDMs and elevated secreted IL-1β and IL-6 levels; these increases were suppressed by hepatocyte-CM derived from *Spp1^f/f^;Alb^Cre^* GAN mice (versus *Spp1^f/^*^f^ GAN) (**Figure 4B,C**). To further verify the relationship between CD44 and TGFBR1 in macrophages, we subjected BMDMs from WT and *Cd44^-/-^* mice to hepatocyte-CM derived from MASH mice. Western blots showed reduced TGFBR1 in *Cd44^-/-^* BMDMs (**Figure 4D**). Given the enrichment of inflammatory genes in *Tgfbr1*^high^ MP1 macrophages (**Figure 3C**), we found a significant reduction in mRNA levels of *Traf6*, *Tlr1*, and *Aim2*, along with reduced ASC (encoded by *Pycard*) in *Cd44^-/-^* BMDMs treated with hepatocyte-CM from MASH mice, compared with WT BMDMs (**Figure 4E,F**). These results suggest that MASH-induced hepatokines prime CD44+ monocytes and activate inflammatory pathways in the transdifferentiated TGFBR1+ macrophages. Consistently, after the GAN diet for 5 months, fewer peripheral CCR2^+^TGFBR1^+^ proinflammatory monocytes and TGFBR1^+^ macrophages infiltrating the atria were detected in *Cd44^-/-^* mice compared with WT (**Figure 4G,H**). While liver phenotype remained unaltered, several atrial arrhythmogenic substrates including structural remodeling and fibrosis were alleviated, and AF susceptibility induced by MASH was reduced in *Cd44^-/-^* mice compared with WT controls (**Figure 4I-M; Supplemental Figure S9, Table S12**), Together, these findings demonstrate that hepatocyte-derived osteopontin activates inflammatory macrophages via CD44 and upregulates TGFBR1 under MASH conditions, underscoring the importance of *Tgfbr1*^high^ MP1 cells in mediating the MASH-driven inflammatory response.

**Figure 4.**
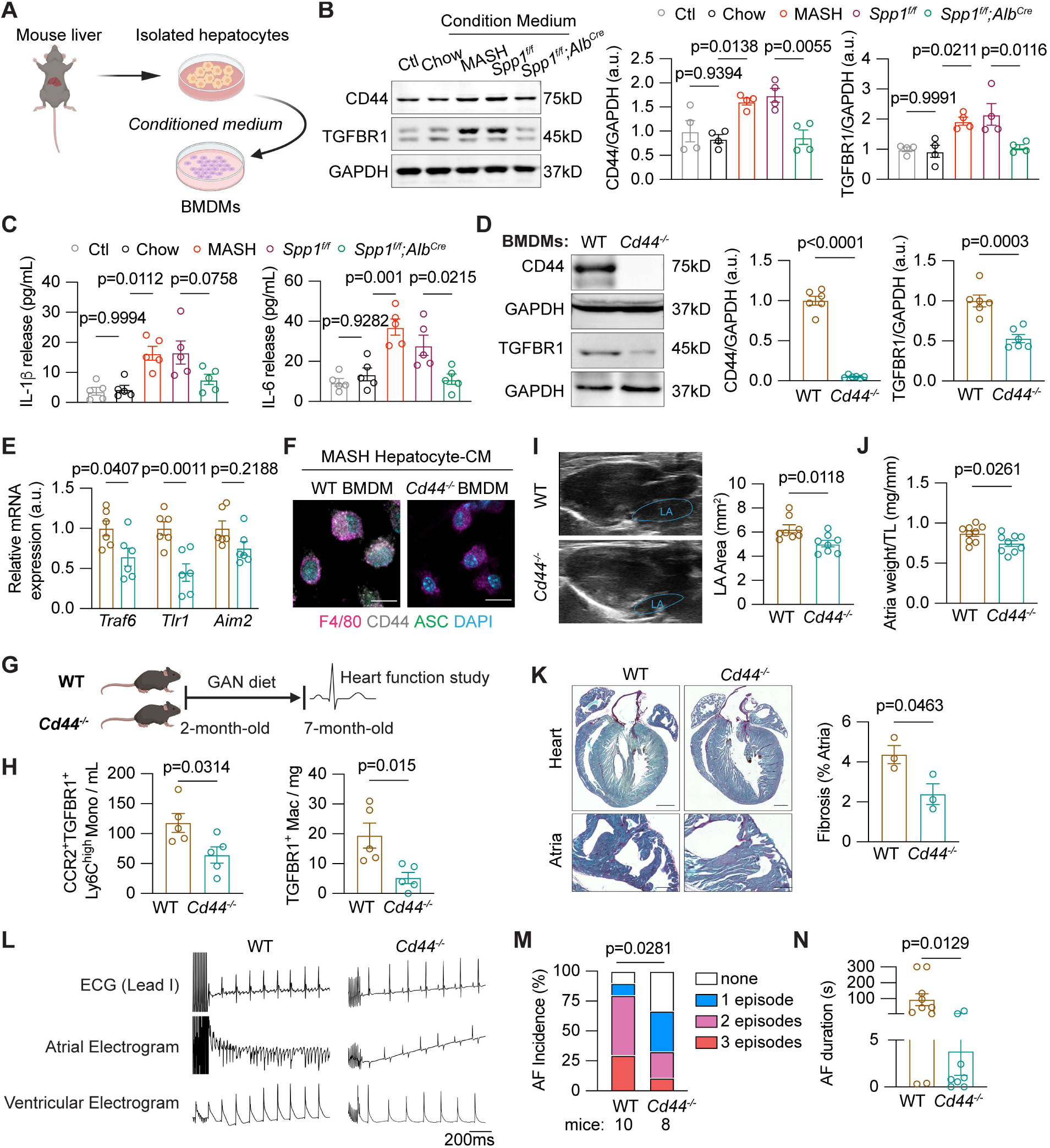
Osteopontin activates *Tgfbr1^high^* macrophages via CD44 receptor. (**A**) Scheme of BMDMs cultured with conditioned medium (CM) from isolated hepatocytes. (**B**) Representative Western blots and quantification of CD44 and TGFBR1 protein in BMDMs cultured in hepatocyte-CM from Chow, MASH, *Spp1^f/f^* GAN and *Spp1^f/f^Alb^Cre^* GAN mice. n=4 per group. (**C**) IL-1β and IL-6 levels secreted from BMDMs cultured with hepatocyte-CM. n=5 per group. (**D**) Representative Western blots and quantification of CD44 and TGFBR1 proteins in WT and *Cd44^- /-^* BMDMs cultured in hepatocyte-CM from MASH mice. n=6 per group. (**E**) Relative mRNA levels of inflammatory genes in BMDMs of WT and *Cd44^-/-^* mice cultured in hepatocyte-CM from MASH mice. n=6 per group. (**F**) Representative images of F4/80, CD44, and ASC in BMDMs of WT and *Cd44^-/-^* mice cultured in hepatocyte-CM from MASH mice. Scale bar 10 μm, n=3 mice per group. (**G**) Scheme of WT and *Cd44^-/-^* mice fed with GAN diet. (**H**) Flow cytometry analysis of peripheral TGFBR1^+^Ly6C^high^ proinflammatory monocytes and atrial TGFBR1^+^ macrophages in WT or *Cd44^- /-^* mice fed with GAN diet. n=5 mice per group. (**I**) Representative B-Mode echocardiography images and measurements of LA areas in WT and *Cd44^-/-^* mice with GAN diet. n=8 per group. (**J**) Quantification of the ratio of atria weight to TL. n=9 per group. (**K**) Representative Picrosirius staining images of four-chamber heart and quantification of atrial fibrosis. n=3 per group. (**L**) Representative surface ECG and intracardiac electrograms of WT and *Cd44^-/-^* mice fed with GAN diet. (**M-N**) Incidence (**M**) and duration (**N**) of pacing-induced AF in two groups. The bar graph data are as mean ± SEM with individual values. p-values are determined with one-way ANOVA Turkey’s multiple comparisons test in **B** and **C**; two-tailed Student’s t-test in **D, E, H, I, J, K** and **N**; Mann-Whitney test in **M.**

### Hepatic osteopontin promotes proarrhythmic multicellular crosstalk in atria

To gain insights into how MASH and hepatic osteopontin changes the cellular landscape of the major non-immune cells within the atria, we characterized the transcriptomic features of three major cell populations – ACMs, FBs, and ECs. GO analysis of the ACMs revealed upregulation of pathways related to “cell junction organization”, “programmed cell death”, and “calcium ion homeostasis” in MASH mice, corroborating the abnormal calcium release events observed in isolated atrial cardiomyocytes. Ablation of hepatic *Spp1* markedly attenuated stress-activated protein kinase signaling cascade and cell migration in ACMs (**Figure 5A; Supplemental Figure S10A**). Consistent with the profibrotic phenotype of MASH atria, FBs showed upregulation of cytokine response, TGFβ signaling, and cellular senescence, which were suppressed by deletion of hepatic *Spp1* (**Figure 5B; Supplemental Figure S10B**). FBs were further classified into 5 subsets (**Figure 5C**). FB1 subset exhibited highest fibroblast activation score (**Figure 5D**), which was selected for subsequent CellChat analysis. ECs exhibited activation of endoplasmic reticulum stress and apoptotic signaling pathways in MASH, which could promote secretory phenotype and leukocyte adhesion; whereas deficiency of *Spp1* diminished cell migration and protein stability in ECs related to cell junction assembly and angiogenesis (**Supplemental Figure S10C,D**). CellChat analysis revealed enhanced FB1-to-FB1 interactions and demonstrated that MP1 exhibited augmented intercellular communications with FB1, ACM, and EC clusters through ITGA, PDGF, IL1B, SIRPB, IGF1, and NPPA pathways, along with increased MP1 autocrine signaling under MASH conditions (**Figure 5E, F**; **Supplemental Figure S10E**). Importantly, ablation of hepatic *Spp1* significantly suppressed the intercellular communications among these populations (**Figure 5E, F**; **Supplemental Figure S10E**).

**Figure 5.**
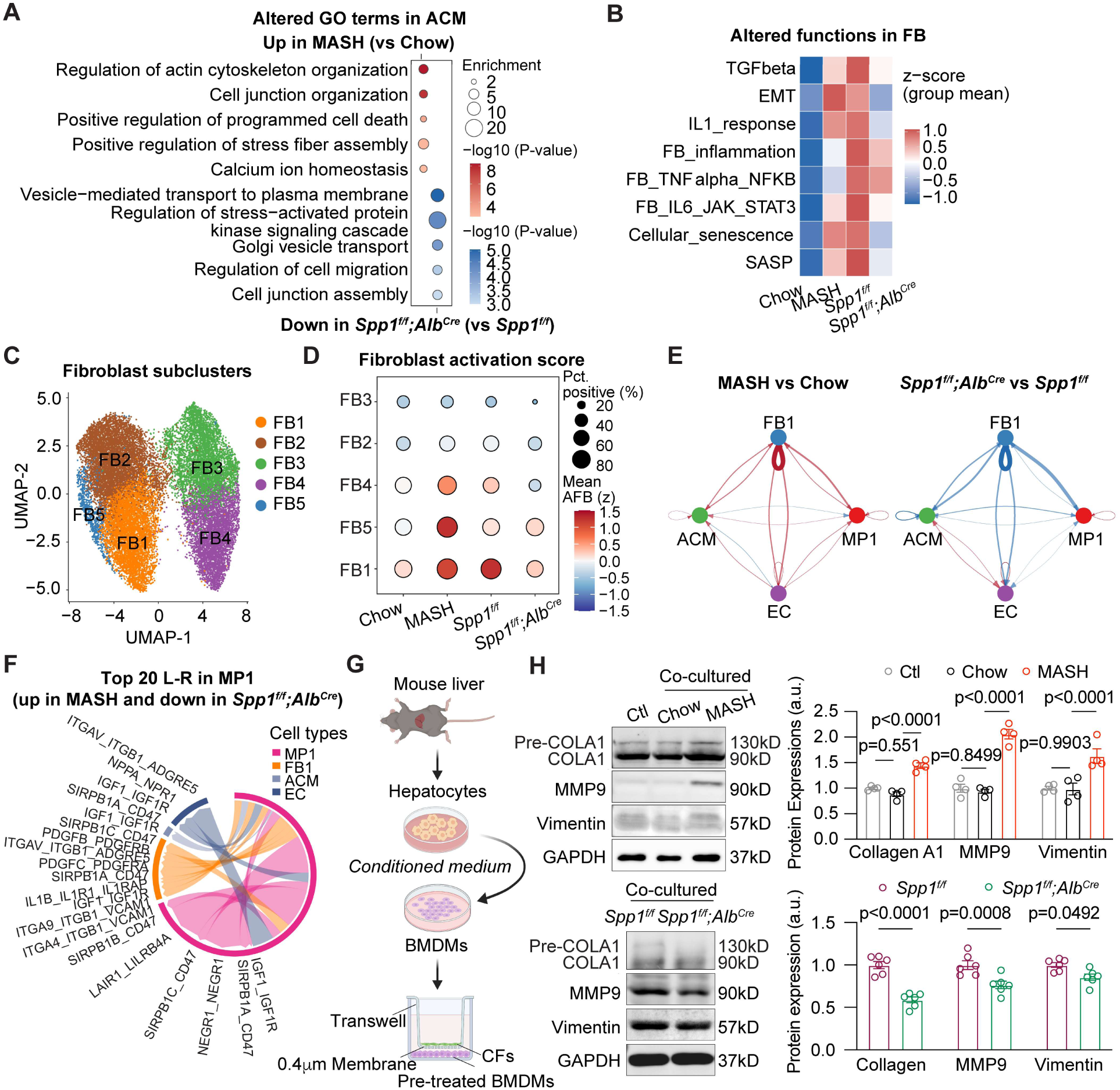
MASH altered macrophage-fibroblast crosstalk through hepatic osteopontin. (**A**) Representative GO terms showing the enhanced pathways in atrial cardiomyocytes (ACM) of MASH mice and the suppressed pathways in ACM of *Spp1^f/f^;Alb^Cre^*mice. (**B**) Representative inflammatory response pathways altered in fibroblast cluster (FB) of mouse atria. (**C**) UMAP visualization of FB subsets. (**D**) Fibroblast activation score across FB subsets in each group. (**E**) CellChat analysis showing differential intercellular interaction strengths among FB1, ACM, EC, and MP1 clusters. (**F**) Top 20 changed ligand-receptor interactions from MP1 to other cell types (EC, FB1, and ACM). (**G**) Scheme of co-culture working system between BMDMs, pre-treated with conditioned medium of isolated hepatocytes from Chow, MASH, *Spp1^f/f^* and *Spp1^f/f^;Alb^Cre^* mice, and adult primary cardiac fibroblasts (CFs) from WT mice. (**H**) Western blots and quantification of Collagen A1, MMP9 and Vimentin proteins in CFs. n=4 or 6 per group. The bar graph data are as mean ± SEM with individual values. p-values are determined with two-tailed Student’s t-test in **H**.

To establish the directionality between inflammatory macrophages and fibroblast activation under the influence of MASH and hepatic OPN signaling, we co-cultured primary adult cardiac fibroblasts (CFs) with BMDMs pre-treated with hepatocyte-CM from MASH mice (**Figure 5G**). CFs co-cultured with BMDMs pre-treated with MASH hepatocyte-CM exhibited increased fibrotic markers (Collagen A1, MMP9, Vimentin) compared with CFs co-cultured with BMDMs pre-treated with chow hepatocyte-CM. In contrast, these increases were attenuated in CFs co-cultured with BMDMs pre-treated with hepatocyte-CM from *Spp1^f/f^;Alb^Cre^* GAN mice (**Figure 5G, H**). Collectively, these findings establish that hepatic osteopontin mediates MASH-induced atrial fibroinflammation by driving the recruitment of inflammatory macrophages, which subsequently activate fibroblasts and impair cardiomyocytes and endothelial cell homeostasis, promoting a arrhythmogenic substrate for AF.

### GSDMD ablation disrupts the liver-to-atria inflammatory axis

Motivated by the upregulation of inflammatory genes in MP1 cluster and the enrichment of programmed cell death and IL-1 pathways of ACM and FB population in MASH atria (**Figure 4C**; **Figure 5A, B**), we next examined the contribution of inflammasome to MASH-induced arrhythmia development. Consistent with the snRNA-seq findings, protein levels of inflammasome components, including cleaved caspase1-p20 and ASC, were elevated in MASH atria, whereas NLRP3 (NACHT, LRR, and PYD domains containing protein 3) and pro-caspase-1 were unchanged between MASH and chow groups (**Supplemental Figure S10F**). As a key downstream effector of the inflammasome pathways, the enhanced cleavage of GSDMD in MASH atria may potentially facilitate the increased secretion of IL-1β and relay fibroinflammatory signal among macrophages and atrial cells. Interestingly, hepatocyte-specific deletion of *Spp1* attenuated the inflammasome activation and GSDMD cleavage in MASH model (**Supplemental Figure S10G**). To directly assess the role of GSDMD in MASH-associated atrial arrhythmia, we subjected 2-month-old GSDMD knockout (*Gsdmd^-/-^*) mice to the GAN diet (**Figure 6A**). GSDMD deficiency reduced inflammasome proteins in MASH atria (**Figure 6B,C**) and suppressed peripheral proinflammatory monocytes and circulating IL-1β and IL-6 levels (**Figure 6D**). Importantly, AF inducibility and duration were reduced in *Gsdmd^-/-^*GAN mice (**Figure 6E-G**), accompanied by reduced atrial weight, normalized atrial size, a trend toward improved left ventricular systolic and diastolic function compared with WT GAN mice (**Figure 6H,I; Supplemental Table S13**). The decreased AF susceptibility in *Gsdmd^-/-^* GAN mice was associated with reduced collagen deposition (**Figure 6J**). Unexpectedly, *Gsdmd^-/-^*mice also exhibited moderately improved liver morphology and function. Liver weight normalized to TL was significantly reduced in *Gsdmd^-/-^* mice compared with WT mice on GAN diet (**Figure 6K**). Although plasma TG, TC, and FPG were comparable between *Gsdmd^-/-^* and WT mice on GAN diet, plasma ALT showed a trend toward reduction (p=0.0717) in *Gsdmd^-/-^*GAN mice, hepatic inflammation and fibrosis were markedly alleviated by GSDMD ablation, whereas steatosis remained unaffected (**Figure 6L-N**). Given that GSDMD executes the activated inflammasome pathway in atrial cardiomyocytes to promote atrial arrhythmia^11^, we employed the cardiomyocytes-specific *Gsdmd* knockout (*Gsdmd^f/f^;Myh6^CreER^*) mice to assess its effect on MASH-induced AF (**Supplemental Figure S11**). The cardiomyocyte-specific deficiency of *Gsdmd* diminished AF incidence and duration and reversed atrial remodeling; these beneficial effects on mitigating AF pathogenesis were independent of hepatic steatosis, inflammation, and fibrosis induced by MASH (**Supplemental Figure S11**). Together, these findings indicate that GSDMD activation in the atrial microenvironment – downstream of osteopontin-mediated macrophage infiltration – is required for atrial arrhythmogenesis in MASH, independent of systemic lipid metabolic alterations.

**Figure 6.**
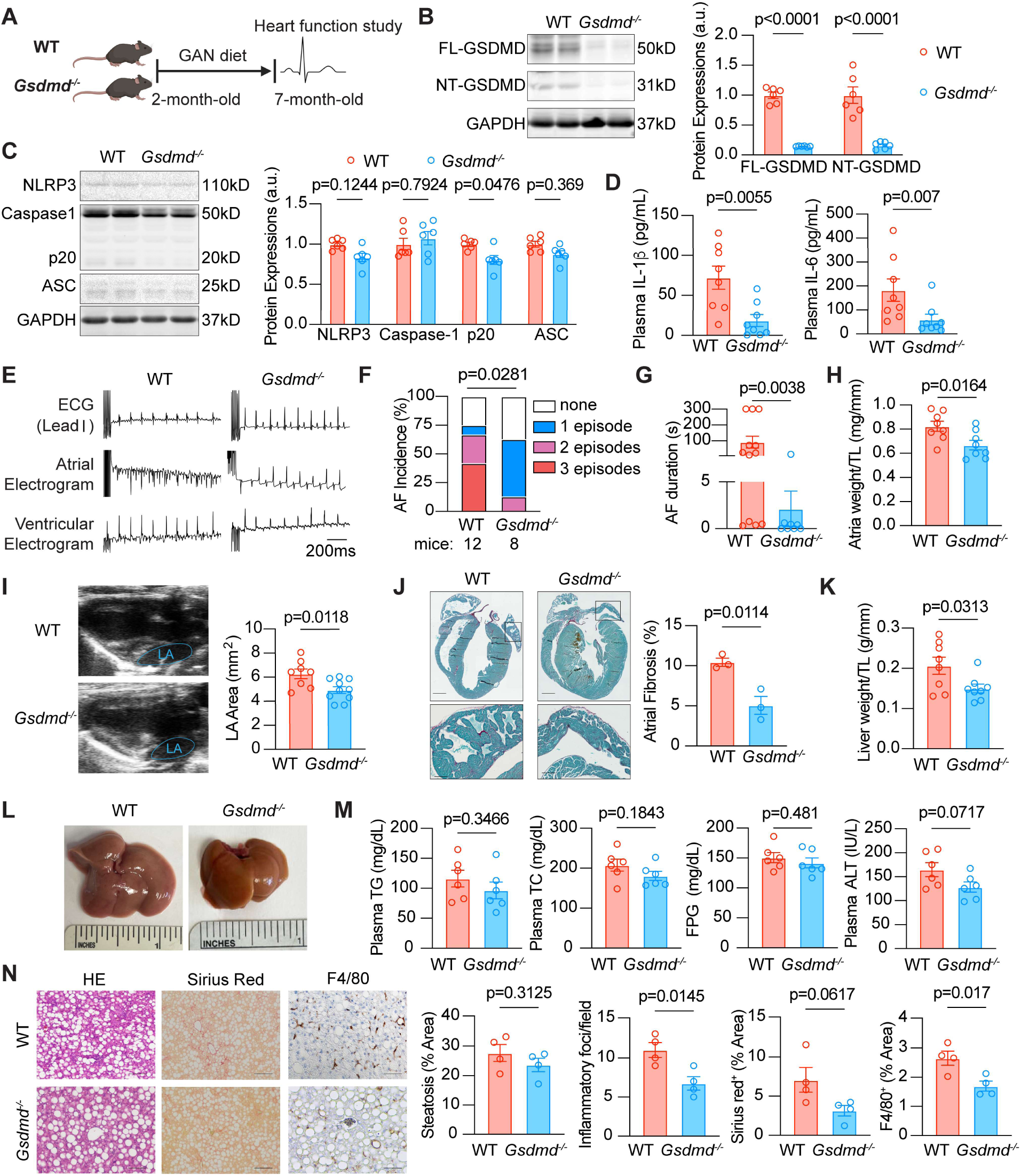
Ablation of GSDMD attenuates MASH-associated AF. (**A**) Schema of 5-month GAN diet feeding in WT and *Gsdmd^-/-^*mice. (**B**) Western blots and quantification of FL-GSDMD and NT-GSDMD proteins in mouse atria. n=6 per group. (**C**) Western blots and quantification of NLRP3, Caspase1, p20 and ASC proteins in mouse atria. n=6 per group. (**D**) Plasma levels of IL-1β and IL-6. n=8 per group. (**E**) Representative surface ECG and intracardiac electrograms in WT or *Gsdmd^-/-^* after 5 months’ GAN feeding. (**F-G**) Incidence (**F**) and duration (**G**) of pacing-induced AF in WT and *Gsdmd^-/-^* mice with GAN diet. (**H**) Quantification of the ratio of atria weight to TL. n=8 per group. (**I**) Representative B-Mode echocardiography images and measurements of LA area in WT GAN and *Gsdmd^-/-^* GAN mice. n=8 or 10 per group. (**J**) Representative Picro-sirius staining images of four-chamber heart and quantification of atrial fibrosis. n=3 per group. (**K**) Quantification of the liver weight to TL ratio in WT GAN and *Gsdmd^-/-^* GAN mice. n=8 per group. (**L**) Representative whole-mount images of livers. (**M**) Plasma levels of TG, TC, FPG, and ALT in two groups. n=6 per group. (**N**) Representative images and quantification of H&E, Sirius red, and F4/80^+^ staining in WT GAN and *Gsdmd^-/-^* GAN mice. n=4 per group. The bar graph data are as mean ± SEM with individual values. p-values are determined with two-tailed Student’s t-test in **B**, **C**, **D**, **E, G, H, I, J, K**, **M**, and **N**; Mann-Whitney test in **F**.

### Neutralizing anti-OPN antibodies ameliorate MASH-associated atrial arrhythmogenesis

Based on dynamically increased OPN levels in MASH mice with longer MASH diet feeding compared with controls (**Figure 7A**), we treated a cohort of MASH mice with either anti-OPN neutralizing antibody (Anti-OPN) or control IgG during the final month of GAN diet regimen (**Figure 7B**). ELISA assays verified that anti-OPN treatment lowered circulating OPN levels and hepatocyte-secreted OPN, without changing body or liver weight (**Figure 7C; Supplemental Figure S12A, B**). Notably, with minimum effect on plasma TG, TC, and FPG levels, anti-OPN antibodies improved hepatic lipid metabolism, steatosis, inflammation, and fibrosis in MASH mice (**Supplemental Figure S12C-G**). Anti-OPN reduced the total number of circulating myeloid cells and Ly6C^high^ monocytes, as well as plasma IL-1β and IL-6 levels compared with the IgG controls (**Supplemental Figure S12H,I**), suggesting that neutralizing osteopontin exerts a systemic anti-inflammatory effect that can mitigate the progression of MASH. Without affecting ventricular function, anti-OPN antibodies reduced atrial size, SCaTs, atrial weight, and atrial collagen deposition in MASH mice, ultimately lowering inducibility and duration of AF in MASH mice (**Figure 7D-J**).

**Figure 7.**
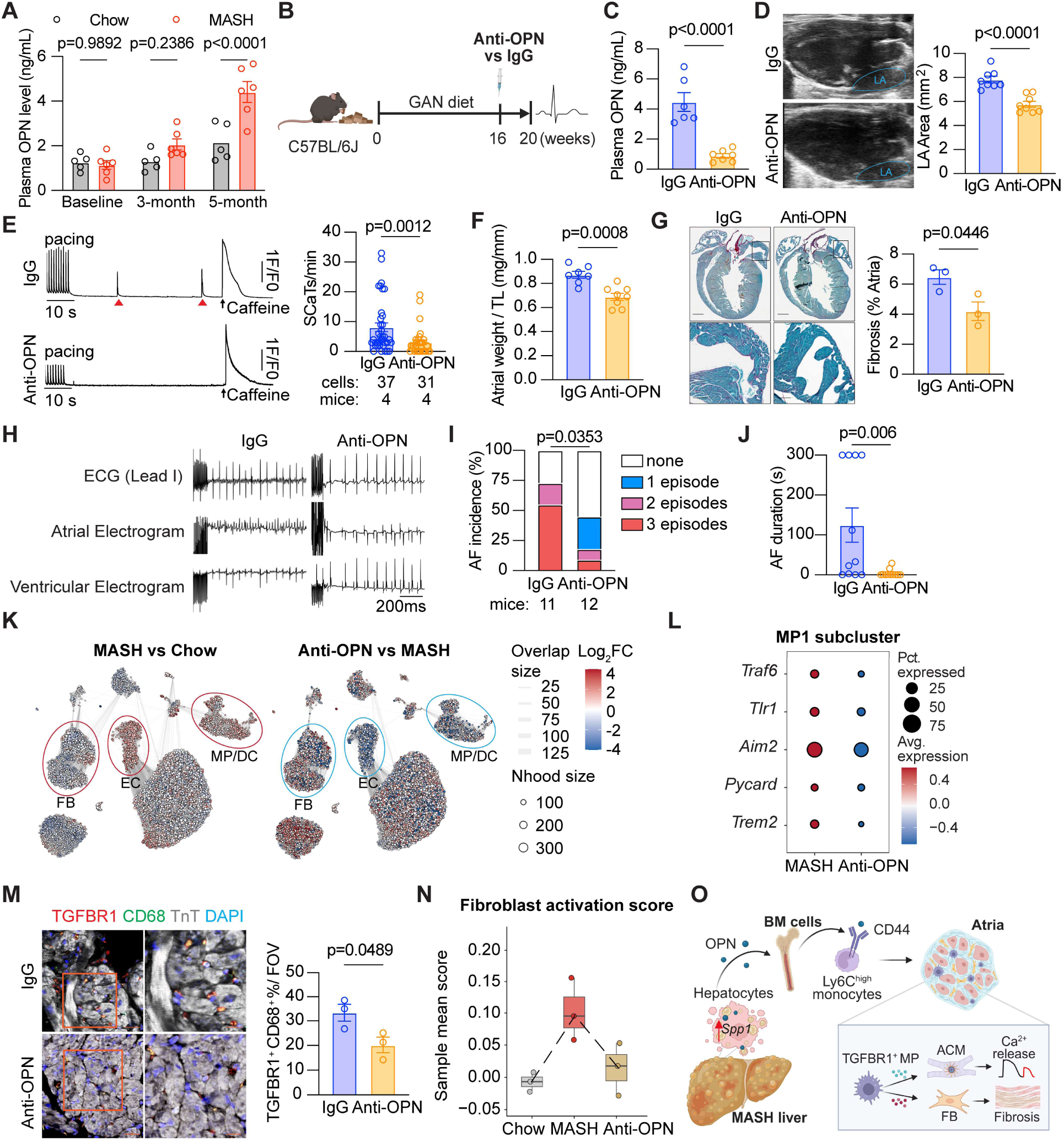
Neutralizing anti-OPN antibody attenuates MASH-associated AF. (**A**) Quantification of plasma OPN levels at baseline, 3 months, and 5 months during chow or GAN diet feeding. n=5 or 6 per group. (**B**) Schema of IgG or anti-OPN antibody treatment in MASH model. (**C**) Plasma OPN levels in IgG- or anti-OPN antibody-treated MASH mice. n=6 or 7 per group. (**D**) Representative B-mode echocardiography images and measurements of LA area. n=8 per group. (**E**) Representative traces of Ca^2+^ transients in isolated atrial cardiomyocytes, followed by baseline recording and the acute application of 10 mmol/L caffeine. Quantification of SCaTs (red arrows). (**F**) Atrial weight to TL ratio. (**G**) Representative Picrosirius staining images of four-chamber heart and quantification of atrial fibrosis, n=3 per group. (**H**) Representative surface ECG and intracardiac electrograms of IgG- and anti-OPN treated MASH mice. (**I-J**) Incidence and duration of pacing-induced AF. (**K**) Milo analysis showing differential cell abundance in each cell population. The MP/DC, FB, and EC clusters were highlighted in circles. (**L**) Dotplot showing downregulated inflammatory genes in MP1 cells in anti-OPN group (vs MASH). (**M**) Representative images of TGFBR1^+^ macrophages in mouse atria. Scale bars, 20 or 10 μm. n=3 mice per group. (**N**) Fibroblast activation scores across 3 groups of FB cluster.. (**O**) The schematic depicts the mechanism by which hepatocyte-derived OPN in MASH promotes multicellular atrial crosstalk through the expansion and recruitment of TGFBR1^+^ inflammatory macrophages, establishing a fibroinflammatory milieu that serves as a proarrhythmic substrate. The bar graph data are mean ± SEM with individual values. p-values are determined with two-tailed Student’s t-test in **A, C, D, E, F, G, J,** and **M**; Mann–Whitney test in **I**.

To examine the mechanism underlying the anti-OPN antibody-mediated anti-arrhythmogenesis, we compared the single nuclei transcriptome between MASH mice and anti-OPN antibody-treated MASH mice. Compared with MASH atria, anti-OPN treatment reduced MP/DCs and ECs and restored ACMs (**Supplemental Figure S13A**). GO terms of pseudo-bulk expression revealed the top upregulated pathways related to regulation of peptidyl threonine phosphorylation, alongside the top downregulated pathways related to IL-6 production and myeloid leukocyte migration in anti-OPN treated atria (**Supplemental Figure S13B**), consistent with the anti-inflammatory response described earlier. Milo plot showed that MASH-induced differentially abundant neighborhoods were reversed by anti-OPN treatment (**Figure 7K**). Specifically, MP/DCs were hyperactivated in MASH, displaying enrichment of inflammatory genes and cytokine production (IL-1 and IL-6), whereas anti-OPN treatment dampened the innate immune response, IL-1β and chemokine production within the MP/DC cluster (**Supplemental Figure 13C**). Proinflammatory genes including *Traf6, Tlr1, Aim2, Pycard, and Trem2* were downregulated in the MP1 subset of anti-OPN treated MASH mice (**Figure 7L**). The infiltration of TGFBR1^+^ inflammatory macrophages into the atria was reduced in anti-OPN-treated MASH mice compared with the IgG-treated MASH mice (**Figure 7M**). Intercellular communications from MP1 to ACM, FB1, and ECs populations were downregulated in pathways related to PDGF, TGFβ, IGF, IL-1β, and CCL signaling (**Supplemental Figure S13D,E**). The increased fibroblast activation score in MASH mice was reduced by anti-OPN treatment (**Figure 7N**). GO analysis revealed that anti-OPN treatment also suppressed functions related to immune response signaling, cell junction organization, and calcium ion import in ACMs, and leukocyte differentiation and angiogenesis in ECs (**Supplemental Figure S13F,G)**. Consistent with the hepatocyte *Spp1* deletion model, anti-OPN treatment also suppressed the protein expressions of the inflammasome pathway in MASH mice (**Supplemental Figure S13H**), underscoring that OPN is an upstream regulator of inflammasome activation. These findings indicate that neutralization of circulating OPN can alleviate the systemic inflammatory response, leading to reduced recruitment of pro-inflammatory *Tgfbr1*^high^ macrophages and a dampened fibroinflammatory milieu in the atria, thereby correcting the proarrhythmic substrate for AF under MASH conditions.

## Discussion

Our work reveals an association between MASH and increased AF risk. With multiple mouse models, we uncover that, under MASH conditions, hepatocyte-derived osteopontin promotes the expansion of proinflammatory TGFBR1^+^ monocytes via the CD44 receptor, which enhances the infiltration of TGFBR1^+^ inflammatory macrophages into the atria, promoting a fibroinflammatory milieu in the atria and Ca^2+^ mishandling conducive to AF development. This liver-to-atria axis can be disrupted through: (i) hepatocyte-specific *Spp1* depletion, which suppresses osteopontin secretion from injured hepatocytes; (ii) anti-osteopontin antibodies, which neutralize the proinflammatory and profibrotic mediator; or (iii) GSDMD ablation, which corrects the response to the enhanced fibroinflammatory milieu (**Figure 7O**).

Emerging evidence suggest that liver dysfunction, such as MASLD/MASH, precedes CVD, and the risk maintains even after adjusting for common comorbidities such as obesity, diabetes, and hypertension^28,29^. Yet, whether MASH is independently associated with AF remains unclear. Using data from the prospective ARIC study, we confirmed that MASH-associated liver fibrosis independently contributes to incident AF over an 8-year follow-up period, as determined by the non-invasive index of liver fibrosis - FIB-4 score ^30^. Given that the FIB-4 index is age-dependent, we utilized the elevated cutoffs of FIB-4 scores for intermediate and advanced livers fibrosis^16^. After adjusting for many well-known AF risk factors including obesity, diabetes, alcohol consumption, total cholesterol, triglycerides, we observed a positive association between incident AF and intermediate and high FIB-4 scores compared with those in the low FIB-4 category. Notably, this association was not significant after further adjustment for prevalent HF, likely because a considerable proportion of patients within the high FIB-4 category were excluded (39 patients, representing 15% of the initial number in this subcategory), rendering the analysis underpowered. In future studies, larger population-based datasets (such as the UK Biobank) should be utilized to increase the sample size within each FIB-4 score category and to validate the observed association between MASH-driven liver fibrosis and incident AF, independent of HF.

A potential concern is whether systemic metabolic dysfunction, rather than liver fibrosis per se, drives AF susceptibility. It is well known that obesity, hyperglycemia, and hypercholesterolemia can independently enhance AF risk. To address this, we adjusted for metabolic factors in Model 2 and Model 3 of the ARIC cohort analysis, which still showed a positive association between incident AF and high FIB-4 index, particularly in high osteopontin level groups. In the mouse studies, the GAN diet-induced MASH model indeed exhibits metabolic syndrome, which could potentially promote the proarrhythmic substrate. However, anti-OPN neutralizing antibody treatment alleviated both the liver phenotype and AF susceptibility in MASH mice without reducing body weight, plasma total cholesterol, or blood glucose levels, suggesting that its protective effect against AF mediates through metabolic status-independent mechanisms. Consistently, both the hepatocyte-specific *Spp1* knockout and Cd44^-/-^ models showed improved liver phenotype and reduced AF incidence, while exhibiting elevated blood glucose and hypercholesterolemia following GAN diet feeding. Taken together, these findings support the notion that hepatic steatosis and fibrosis directly promote atrial arrhythmogenesis through key hepatokines, such as osteopontin, independent of hyperglycemia and hypercholesterolemia.

To validate the clinical findings, we employed a classical preclinical GAN diet induced MASH model, which recapitulates the characteristics of human MASH including steatosis, inflammation, and fibrosis. Mice fed the GAN diet exhibited markedly increased susceptibility to AF compared with the chow-fed controls, without elevation in blood pressure or worsening in exercise endurance. Echocardiography revealed preserved left-ventricular diastolic function, excluding concurrent heart failure with preserved ejection fraction (HFpEF) phenotype. Consistently, another study reported that MASH-induced *foz/foz* mice developed compensated hypertrophy, and only exogenous angiotensin II amplified decompensated diastolic function^31^. Although epidemiological data suggest that MASLD and MASH are independently associated with HFpEF ^32–34^, whether MASH predisposes to HFpEF independent of AF burden remains unknown. Our finding suggest that AF may precede HFpEF by promoting hemodynamic congestion and diastolic impairment under MASH condition. Long-term GAN-diet models with serial assessment of atrial electrophysiology and ventricular filing dynamics will be helpful to clarify the temporal relationship between AF and HFpEF progression during MASH.

Inflammation and fibrosis are important mediators of MASH and AF pathogenesis^35,36^. Innate immune cells display substantial heterogeneity in origin and function, amplifying hepatic and systemic inflammation in MASH^37,38^. However, no study has characterized the immune cell signatures within MASH atria or their direct impact on atrial remodeling^39^. Our snRNA-seq revealed the cellular landscape of MASH atria, showing increased recruitment of the MP/DC population, reduced cardiac muscle contraction, disruption of calcium homeostasis, and activation of inflammasome/GSDMD pathways. In particular, under MASH conditions, a subset of monocyte-derived macrophages with high expression of TGFBR1 became more proliferative and proinflammatory, driving pyroptosis and IL-1 production. Given our recent findings demonstrating a role of cardiomyocyte GSDMD activation in promoting atrial remodeling^11^, we now show that global ablation of *Gsdmd* not only reduced infiltration of TGFBR1^+^ macrophages, IL-1β production, and atria fibrosis, but also moderately improved hepatic inflammation and fibrosis, while steatosis remained unaffected. Furthermore, the cardiomyocyte-specific *Gsdmd* deletion only attenuated MASH-induced AF susceptibility but did not alter liver phenotype. These data suggests that hepatic GSDMD signaling may be involved in liver fibrosis, which requires further investigation. Nevertheless, these findings also position GSDMD as a downstream effector of the MASH-induced hepatokines such as osteopontin and the associated systemic inflammatory response.

Hepatokines emerge as critical mediators of liver-heart communication and affect CVD progression^21^. Consistent with prior studies^40,41^, our proteomic profiling revealed elevated circulating osteopontin levels in MASH mice. We identified that hepatocytes, rather than liver resident macrophages, are the primary source of osteopontin within the liver. Osteopontin exerts pleiotropic and context-dependent effects in modulating immune-cell reprogramming via its receptors CD44 and integrin β1 in MASLD and MASH condition ^42,43^. On one hand, osteopontin facilitates the expansion and migration of monocyte-derived macrophages into local lesion to amplify the inflammatory response^44^; on the other hand, depletion of *Spp1* in lipid-associated or resident hepatic macrophages exacerbates MASH progression^45^. Both genetic ablation and pharmacologic inhibition of osteopontin have been shown to attenuate hepatic inflammation and fibrosis ^46,47^. Here, we demonstrate that hepatocyte-specific *Spp1* ablation mitigates both liver dysfunction and pro-arrhythmic atrial remodeling by suppressing the activation of proinflammatory TGFBR1+ macrophages. Consistently, blocking the osteopontin receptor CD44 also attenuated MASH-induced proinflammatory remodeling and the proarrhythmic substrate in the atria. These data establish the hepatocyte SPP1 – CD44+TGFBR1+ macrophage axis in MASH-induced arrhythmogenesis. A recent study has demonstrated the cell-type specific silencing of *Spp1* via antibody-siRNA conjugates alleviates proarrhythmic substrate such as fibrosis in the atria^48^. Together with our findings on anti-OPN antibodies, this evidence positions osteopontin as a novel target for AF. The benefits of the anti-OPN approach are accompanied by reduced recruitment of inflammatory macrophages into MASH atria. In human cohorts, higher plasma OPN levels were correlated with increased AF risk among individuals with elevated FIB-4 scores, suggesting its potential as a biomarker for advanced liver fibrosis and atrial remodeling. Of note, our proteomics data demonstrated increases in several hepatokines under MASH conditions, the roles of which in AF pathogenesis have not been studied. Future studies are needed to determine whether other hepatokines play a role in MASH-induced systemic immune responses as well as MASH-related cardiac pathology.

Our study has several limitations. First, our observational human study only revealed the association but not causation between liver fibrosis severity and AF; prospective studies are warranted to confirm whether elevated circulating osteopontin levels serve as a reliable biomarker for liver fibrosis and MASH-associated AF. Second, the upregulation of *SPP1* was identified in a subset of hepatocytes based on a public snRNA dataset from MASH patients; whether SPP1 upregulation in human livers or hepatocytes is directly associated with AF occurrence in MASH patients remains to be determined. Third, while the GAN-diet induced MASH model recapitulates key characteristics of hepatic fibroinflammation, validation in a large animal model of MASH would further strengthen the translational relevance of our findings. Finally, while we identified hepatocytes as an important resource of osteopontin under MASH conditions, the contribution of osteopontin secreted by other cell types or organs to the inflammatory response and AF development cannot be excluded.

## Conclusions

This study demonstrates that MASH promotes atrial arrhythmogenesis through the hepatocyte-derived osteopontin. This liver-to-atria signaling circuit enhances inflammatory macrophage recruitment into the atria, creating a fibroinflammatory milieu that favors the development of AF substrate. Interventions such silencing hepatic *SPP1*, neutralizing osteopontin, or blocking inflammasome effectors offer therapeutic potential for mitigating MASH and its related arrhythmogenesis.

## Author contributions

Y. Yuan, B. Dong, and N. Li designed this study. Y. Yuan, S. Wang, Y. Zeng, T. Li, A. K. Shinohara, C Lin, and S. Y. Jung conducted the experiments. J. Ding and J. Jiang performed the bioinformatics analysis. D. Olivares-Villagómez provided *Spp1* flox mice. Y. Yuan, S. Wang, C. W. Sun, R. C.Hoogeveen, and S. Saadatagah analyzed the data. B. Dong, C.M. Ballantyne, and N. Li conceptualized the study. M. G. Chelu, C. M. Ballantyne, B. Dong, and N. Li supervised the research. Y. Yuan and N. Li wrote the original draft. S. Wang, S. Saadatagah, C.M. Ballantyne B. Dong, and N. Li reviewed and edited the manuscript.

## Abbreviations and acronyms

ACM: atrial cardiomyocyte
AF: atrial fibrillation
ALT: alanine aminotransferase
AST: aspartate aminotransferase
BM: bone marrow
BMDM: bone marrow derived macrophage
CM: conditioned medium
ECG: electrocardiogram
FPG: fasting plasma glucose
GAN: Gubra-Amylin NASH
GSDMD: gasdermin D
HFpEF: heart failure with preserved ejection fraction
MASH: metabolic dysfunction–associated steatohepatitis
MASLD: metabolic dysfunction-associated steatotic liver disease
NLRP3: NLR family pyrin domain containing 3
SCaTs: spontaneous Ca2+ transients
OPN: osteopontin
TC: total cholesterol
TG: triglyceride

